# Discordant evolution of mitochondrial and nuclear yeast genomes at population level

**DOI:** 10.1101/855858

**Authors:** Matteo De Chiara, Anne Friedrich, Benjamin Barré, Michael Breitenbach, Joseph Schacherer, Gianni Liti

## Abstract

Mitochondria are essential organelles partially regulated by their own genomes. The mitochondrial genome maintenance and inheritance differ from nuclear genome, potentially uncoupling their evolutionary trajectories. Here, we analysed mitochondrial sequences obtained from the 1,011 *Saccharomyces cerevisiae* strain collection and identified pronounced differences with their nuclear genome counterparts. In contrast with most fungal species, *S. cerevisiae* mitochondrial genomes show higher genetic diversity compared to the nuclear genomes. Strikingly, mitochondrial genomes appear to be highly admixed, resulting in a complex interconnected phylogeny with weak grouping of isolates, whereas interspecies introgressions are very rare. Complete genome assemblies revealed that structural rearrangements are nearly absent with rare inversions detected. We tracked introns variation in *COX1* and *COB* to infer gain and loss events throughout the species evolutionary history. Mitochondrial genome copy number is connected with the nuclear genome and linearly scale up with ploidy. We observed rare cases of naturally occurring mitochondrial DNA loss, *petite*, with a subset of them that do not suffer fitness growth defects. Overall, our results illustrate how differences in the biology of two genomes coexisting in the same cells can lead to discordant evolutionary histories.

## Introduction

Mitochondria are pivotal in eukaryotic cells, providing chemical energy in the form of ATP through oxidative phosphorylation (Saraste 1999), but also fulfilling a multitude of other critical functions including lipid and amino acids metabolism as well as heme and nucleotide synthesis (Spinelli and Haigis 2018). These fundamental organelles share a common origin in eukaryotes deriving from ancestral α-proteobacterial symbiont (Gray et al. 1999) and are still present in the vast majority of eukaryotes (Tovar et al. 1999; Karnkowska et al. 2016). Since then, mitochondrial genomes have undergone an drastic size reduction by gene transfer to the nucleus (Wallace 2007; Gray 2012), although they evolved in a large variety of sizes and gene content across species (Burger et al. 2013; Flegontov et al. 2015). Despite having their own genomes and translation machinery, mitochondria nevertheless remain strongly interconnected with the host cell. Several proteins are encoded at nuclear level but they are relocalized into the mitochondria once mature. For example, eukaryotic ATP synthase subunits are encoded by both nuclear and mitochondrial genomes. Consequently, the products of nuclear encoded subunits need to be imported into the organelles and assembled in a coordinate manner into functioning structures (Rühle and Leister 2015). The tight interplay between genes coded by nuclear and mitochondrial genomes has thus driven the coevolution of these interacting proteins and incompatibilities between mitochondrial and nuclear alleles may contribute to reproductive barriers between species (Lee et al. 2008) or subspecies (Ma et al. 2016).

The mitochondrial DNA (mtDNA) has been widely used to infer phylogenetic relationships among species and individuals (Atyame et al. 2011; Lassiter et al. 2015; Nadimi et al. 2016; Turissini et al. 2017). However, mtDNA phylogenies can vary from those generated from nuclear genome (Fontenot et al. 2011). Mitochondrial genome is usually found in high copy number with multiple copies in each organelle (Wiesner et al. 1992). Mutation rate of mtDNA is also generally different from the one of the nuclear DNA and it can be significantly lower (plants and fungi) or higher (animals) (Sandor et al. 2018). In addition, mtDNA does not follow a mendelian inheritance, since its replication and partition is not directly linked to the cell cycle and transmission is typically uniparental (maternal, in the vast majority of cases), eliminating the potential for sexual recombination. Cases of paternal or biparental inheritance have nevertheless been reported in some species (Zouros et al. 1992; Leducq et al. 2017), although mtDNA heterogeneity does not persist. For instance, as in *Mytilidae* mussel species, where paternal mitochondria are only transmitted to the gonads of male offspring (Zouros et al. 1994) and in ascomycetes yeasts mitochondrial heteroplasmy is lost after the first few mitotic cycles (Solieri 2010).

Given the importance of mitochondrial functions, deleterious mutations often cause severe and untreatable diseases (Schapira 2012). Given the difficulty to engineer mtDNA in human cell lines, model genetic systems have paved the way for functional characterization. In particular, the budding yeast *S. cerevisiae*, has been widely used to study the phenotypic effect of mtDNA variation (Ghosh et al. 2014; Lasserre et al. 2015; Song et al. 2016). *S. cerevisiae* mtDNA is very large, 85 kb, and organized as either circular monomer or head-to-tails tandem-repeated linear structures (Solieri 2010). The *S. cerevisiae* mtDNA contains eight genes encoding for three subunits of the ATP synthase complex (*ATP6*, *ATP8*, *ATP9/OLI1*), the apocytochrome b (*COB*), three subunits of the cytochrome c oxidase complex (*COX1*, *COX2*, *COX3*) and a ribosomal protein (*VAR1/RPS3*). Recombination can occur between yeast mitochondria during the transient heteroplasmic phase following the mating. However the two parental organelles are kept physically separated and can only be in contact and recombine on a limited interaction surface (Fritsch et al. 2014). The S288C laboratory genetic background contains approximately 20 copies of mitochondrial genomes and 5% of non-essential genes are required for mtDNA maintenance (Puddu et al. 2019). Cells with mutated or without mitochondrial genomes are referred as “petite” given their slow growth phenotype in complete media and are unable to grow on non-fermentable carbon sources. Although *S. cerevisiae* is a leading model for molecular and genomics studies (Tsukada and Ohsumi 1993; Botstein and Fink 2011; Cáp et al. 2012), most studies have focused on a small number of laboratory derivative isolates, whose genetic and phenotypic features have been shaped by laboratory manipulations (Warringer et al. 2011). Indeed, high mitochondrial genome instability, which promote petite formation, is a hallmark of *S. cerevisiae* laboratory strains (Dimitrov et al. 2009).

The release of mitochondrial genomes of several isolates (Bergström et al. 2014; Strope et al. 2015) offered an opportunity for mitochondrial population surveys (Wolters et al. 2015). We recently released high coverage genome sequence for a panel of 1,011 natural strains, isolated from both anthropic and wild niches (Peter et al. 2018). The nuclear genome phylogeny revealed strong population stratification with 26 separated lineages and three groups of outbred mosaic strains. These lineages were further partitioned in domesticated and wild lineages based on the ecological origin of the isolates. Nuclear genome diversity is different between these two classes of isolates, with wild lineages having a higher number of SNPs, while domesticated showing higher genome content variation. Here, we examined mitochondrial genome variation across the 1,011 *S. cerevisiae* collection to trace the events that shaped their evolution and compare them with evolutionary patterns observed in the nuclear genomes. We detected strong admixture of the mtDNA, which cannot be fully captured by phylogenetic trees. The mitochondrial genomes have higher genetic variation compared to their nuclear genome counterparts with weak concordance between the two phylogenetic topologies. In addition to sequence variability, we observed high variation in mtDNA copy number and identified several natural petite isolates, with some that surprisingly fully recovered growth fitness defects.

## Results and Discussion

### Mitochondrial genetic diversity across *S. cerevisiae* population

We explored 1,011 *S. cerevisiae* sequenced isolates (Peter et al. 2018) to investigate the intraspecific mitochondrial genome diversity and evolution. Since mitochondrial genomes include long AT-rich and variable intergenic regions that are difficult to compare, we first focused on the eight mitochondrial coding DNA sequences (CDSs). From 698 *de novo* genome assemblies, we collected the eight complete CDSs. Out of these, 553 isolates also had complete or nearly complete mitochondrial sequence. A subset of 353 genome sequences did not have any ambiguous base across the CDSs (supplementary fig. S1, supplementary table S1). We estimated the global genetic diversity by the average pairwise divergence π. Overall, we observed lower diversity in the coding nuclear (π ~0.003) (Peter et al. 2018) compared to the mitochondrial sequences (π ~0.0085, supplementary table S2), which contrasts to what was previously observed for other yeast species (supplementary fig. S2). This trend is more similar to the pattern observed in animal rather than in fungi (Freel et al. 2014; Sandor et al. 2018).

We observed sharp genetic divergence differences of the nuclear and mitochondrial genomes among wild and domesticated isolates. In wild clades, despite higher nuclear divergence (up to 1.1% at CDS level), the mitochondrial CDS genetic distance reaches its maximum of ~0.6% at ~0.4% of nuclear divergence and plateau afterwards. In contrast, mitochondrial sequence divergence between domesticated clades have a larger increase, reaching its maximum at lower nuclear divergences (supplementary fig. S3). This difference in variation is observed across all the mitochondrial CDSs whose values of π are systematically higher in domesticated compared to wild isolates.

Shortest CDSs, *ATP8* and *ATP9*, have the lowest proportion of polymorphic sites (~2%), lowest values of π (3E-03 or less) and lack of non-synonymous mutations. In contrast, *COX1* and *COX2* are highly polymorphic. Although *COX1* has the highest polymorphic sites (8%), *COX2* has the highest π value (0.0163, table 1). We used the Discriminant Analyses of Principal Component (DAPC) (Jombart et al. 2010) to evaluate the contribution of specific genes to classify mitochondrial ‘haplotypes’ and population clustering. We quantified that *ATP6* and *COX2* respectively account for 38% and 28% of population clustering. This observation supports the widespread usage of *COX2* in mitochondrial phylogeny (fig. 1a) (Kurtzman and Robnett 2003; Peris et al. 2014; Peris et al. 2016; Peris et al. 2017).

**TABLE 1.** Genetic diversity of mitochondrial CDSs

|  | N. of<br>complete<br>CDSs | Len. of<br>CDS in<br>S288c | Len. of<br>alignment | N. of<br>polymorphic<br>sites | InDel<br>positions | pi | dN/dS<br>(median<br>value) |
| --- | --- | --- | --- | --- | --- | --- | --- |
| **ATP6** | 933 | 780 | 780 | 41 | 0 | 0.0108 | 0.1503 |
| **ATP8** | 971 | 147 | 147 | 3 | 0 | 0.0034 | 0 |
| **ATP9** | 951 | 231 | 231 | 4 | 0 | 0.0013 | 0 |
| **COB** | 774 | 1158 | 1158 | 44 | 0 | 0.0048 | 0.0753 |
| **COX1** | 737 | 1605 | 1605 | 129 | 0 | 0.0088 | 0.1034 |
| **COX2** | 909 | 756 | 756 | 45 | 0 | 0.0163 | 0.0288 |
| **COX3** | 935 | 810 | 810 | 36 | 0 | 0.0072 | 0.0222 |
| **VAR1** | 453 | 1197 | 1302 | 82 | 190 | 0.0070 | 0.2500 |

**FIG. 1.**
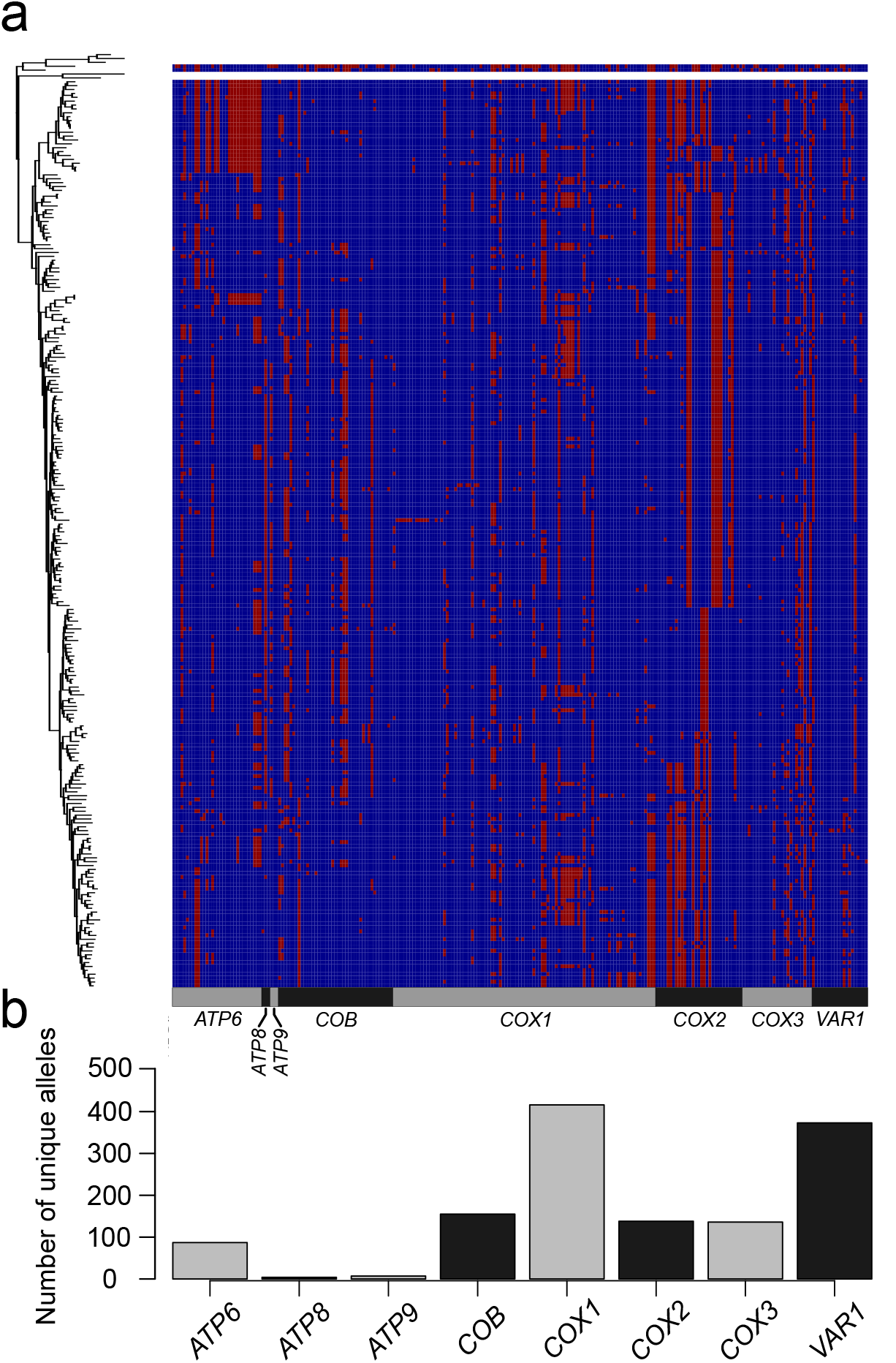
Allele distribution across mitochondrial CDSs. (a) Distribution of major (blue) and minor (red) alleles for the 259 polymorphic positions in the 234 *S. cerevisiae* unique complete profiles (which include 353 isolates). Profiles are ordered according to their phylogenetic relationship using the phylogenetic Neighbour-Joining tree (left side). (b) The numbers of unique alleles for each mitochondrial CDS show dramatic difference between genes.

Next, we generated a non-redundant allele database. We observed a variable number of distinct CDSs alleles (fig. 1b), resulting in high proportion unique allelic profiles (234 out of 353 isolates, supplementary fig. S1). We used these non-redundant allelic profiles as a proxy to investigate the mitochondrial genome distribution in the population. Globally, we observed a poor overlap between the mitochondrial and nuclear genome phylogenetic lineages (Peter et al. 2018) with few exceptions that include near-to-clonal nuclear genome lineages, having specific mitochondrial profiles. These exceptions include a Sake subclade, the two clinical Wine/European subclades (Y’ amplification and *S. boulardii*), the North American and the reproductively isolated Malaysian clades (Liti et al. 2009). In contrast, the Mixed Origin clade (Peter et al. 2018), which has highly diverse ecological (*e.g*. bakeries, beer, plants, animal, water, clinical sample) and geographical origins (*e.g*. Europe, Asia, Middle East, America), shows low mitochondrial intra-clade difference despite substantial nuclear genome variation (supplementary fig. S4). Indeed, across the Mixed Origin clade, only very similar profiles of mitochondrial genes segregate, with variants limited to *COX1* and *VAR1*, resulting in very low π (8.0E-05) compared to other clades (~E-03, supplementary table S2).

The *VAR1* gene is a particularly variable, highly AT rich and prone to non-synonymous mutations and indels. These indels mostly consist of GC clusters that are byp-like elements causing jumps in protein translation in other yeast species (Lang et al. 2014). Two positions were described, one named ‘common’ and another downstream with GC cluster in inverted orientation (Nosek et al. 2015). We identified 35 allelic variants of *VAR1* gene harbouring these two clusters in 117 isolates, mainly belonging to the mosaic groups (N=52) (supplementary table S1). While most of the reported cases harboured the GC cluster either in the common (N=91, accounting for 18 different *VAR1* alleles) or in both positions (N=6, across 4 *VAR1* alleles) (Nosek et al. 2015), a large fraction of the observed allelic variants here only harboured the GC cluster in the second site (N=19, across 13 *VAR1* alleles). We also discovered two novel variants, one with GC cluster at the common position but in inverted orientation (2 isolates) and a second with the GC cluster in tandem duplication at the second position (3 isolates). Altogether, these results uncovered high variability of mitochondrial sequence across the *S. cerevisiae* natural population.

### Extensive admixture of mitochondrial genomes

We investigated the mitochondrial genome population structure using the eight concatenated CDS to calculate the phylogenetic network using SPLITSTREE (Huson and Bryant 2006). The dataset comprises 239 non-redundant CDS profiles, limited to the 234 *S. cerevisiae* isolates with the complete set of CDS sequences, and included the five representatives of *S. paradoxus* clades (Yue et al. 2017) as out-groups. The resulting intertwined network shows a strong interconnectivity of the sequences, underlying frequent historical recombination (fig. 2a). In contrast, classical phylogenetic trees are unable to consistently group the isolates (fig 2b). Using ADMIXTURE (Alexander et al. 2009), we observed that the opposite edges of the trees fall in the same population for low K values (K=2-3), further underlying a poor grouping.

**FIG. 2.**
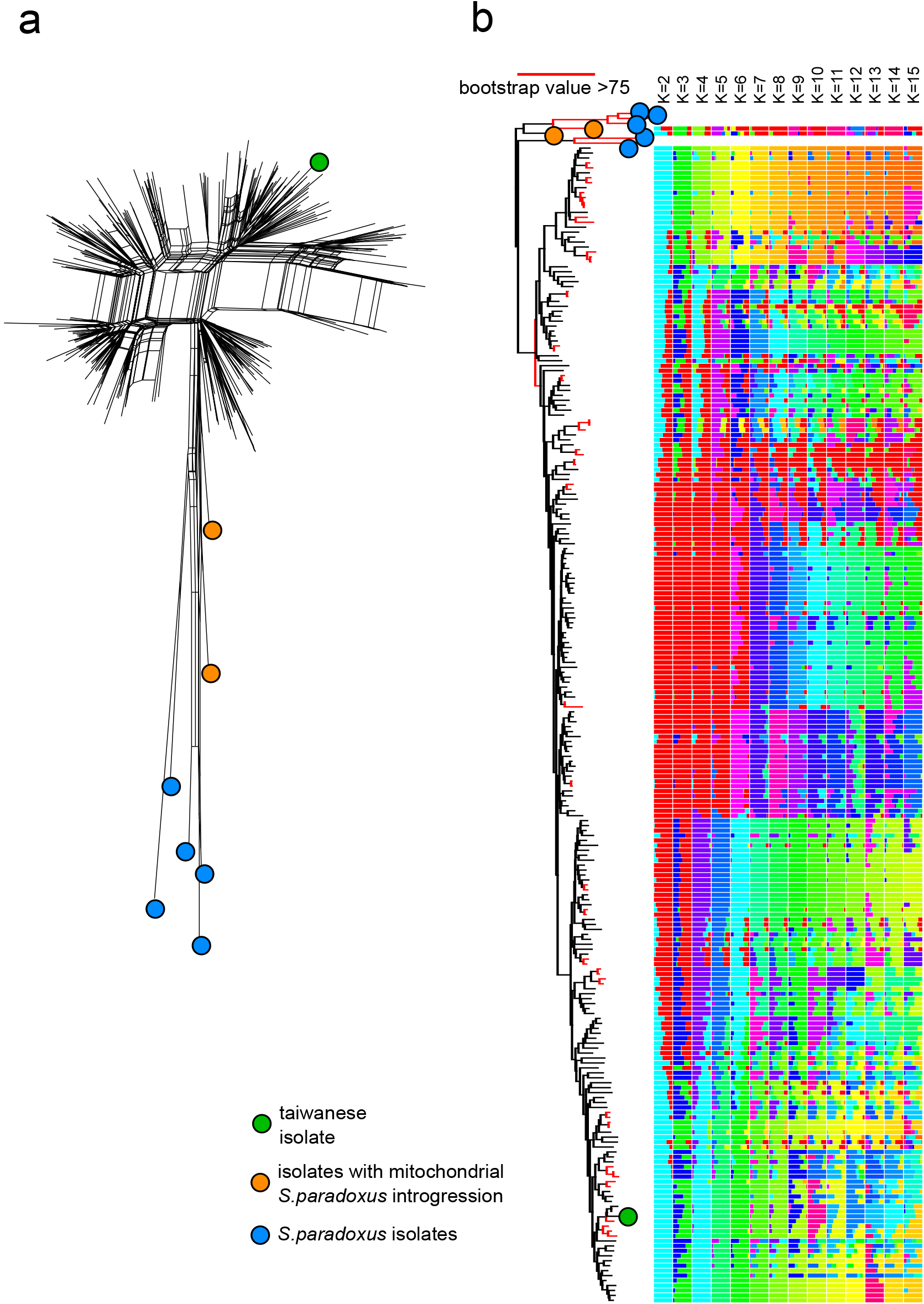
Complex mitochondrial genomes phylogeny. Only *S. cerevisiae* isolates with complete CDS data have been used (N=353) in addition to five *S. paradoxus* isolates used as outgroup. (a) Phylogenetic network of non-redundant concatenated CDS sequences (N=237 profiles) produced a highly intertwined network driven by recombination with few groups of closely related strains. (b) The rooted tree (left) shows a weak topology with few nodes (red) with bootstrap values over 75. ADMIXTURE analysis of genomic components (right) with K ranging from 2 to 15 confirms the high degree of mosaicism. The highly divergent Taiwanese lineage (green dot) is not basal to the other lineages in contrast to the nuclear genome phylogeny.

Mitochondrial population structure appears to poorly reflect the clustering obtained from the nuclear genome. For example, the most divergent and basal Taiwanese clade based on nuclear genome, does not show a higher mitochondrial diversity. However, isolates belonging to the Mosaic groups of the *S. cerevisiae* population show the highest degrees of admixture, indicating that outbreeding has impacted both mitochondrial and nuclear genomes. We then calculated the coefficient of concordance ‘W’ using the CADM (Congruence Among Distance Matrix) metric (Campbell et al. 2011) to be 0.79, with 0 indicating complete disagreement and 1 complete agreement between distance matrixes. This value indicates a relatively good concordance between the phylogenic networks of mitochondrial and nuclear genomes. This is likely driven by isolates with very close mitochondrial sequence often also having similar nuclear genome sequence, while the main branches of the mitochondrial tree cannot be positioned with confidence. Overall, our results highlight a pronounced separation in evolutionary histories of the two co-existing genomes and the extensive recombination found provides additional support to yeast mitochondrial inheritance requiring recombination-driven replication (Kowalczykowski 2000; Chen and Clark-Walker 2018; Dujon 2019).

### Interspecies introgressions of mtDNA are rare

We recently described four clades (namely Alpechin, Mexican Agave, French Guiana and Brazilian Bioethanol) with abundant *S. paradoxus* interspecies introgressions in the nuclear genome (Peter et al. 2018). We analysed the mitochondrial CDSs to search for introgressed alleles. The four clades with abundant nuclear genome introgressions did not show any *S. paradoxus* mitochondrial alleles. Nevertheless, two isolates from America (CQS, YCL) and one from Africa (ADE), all genetically related to the French Guiana and the Mexican Agave clades, harbour two distinct patters of *S. paradoxus* mitochondrial introgressions. We could retrieve the complete CDSs set for two of them (CQS and YCL), while the third (ADE) is incomplete but very close to YCL. The mitochondria introgression in YCL (YJM1399) strain was already reported, but no further analyses were presented (Wolters et al. 2015). We generated a set of polymorphic markers (methods), to accurately identify the introgression boundaries. The *S. cerevisiae* major alleles were identified from the 1,011 isolates, whereas for *S. paradoxus* we derived a consensus sequence from three high quality sequences encompassing the North and South American as well as Hawaiian variation (Yue et al. 2017). The European and Far East Asian *S. paradoxus* isolates were not included because of their similarity with *S. cerevisiae* sequences, likely due to an ancient introgression event from *S. cerevisiae* to *S. paradoxus* (Wu and Hao 2015; Leducq et al. 2017; Yue et al. 2017). We generated a catalogue of 116 polymorphic positions and derived different alleles between the two species. Several genes in these two isolates were catalogued either as partially or fully introgressed (fig. 3a). Since the frequency of some alleles is close to 50% and often the less common allele of one species is the more common allele of the second one, there is a chance of calling false positive introgressions. Nevertheless, long consecutive series of *S. paradoxus* marker in the *COB*, *ATP9*, *COX1*, *COX2* and *COX3* genes in YCL, as well as those in the *COB*, *COX1*, *COX2* and *COX3* genes in CQS are likely to be genuine. The absence of traces of introgression in *S. cerevisiae* isolates from Europe could be explained by the higher sequence similarity with European *S. paradoxus*, which prevent the detection. However, introgressions between *S. cerevisiae* and European *S. paradoxus* isolates could also be prevented by the non-collinearity in the structure of their mitochondrial genomes that likely impair recombination (Yue et al. 2017).

**FIG. 3.**
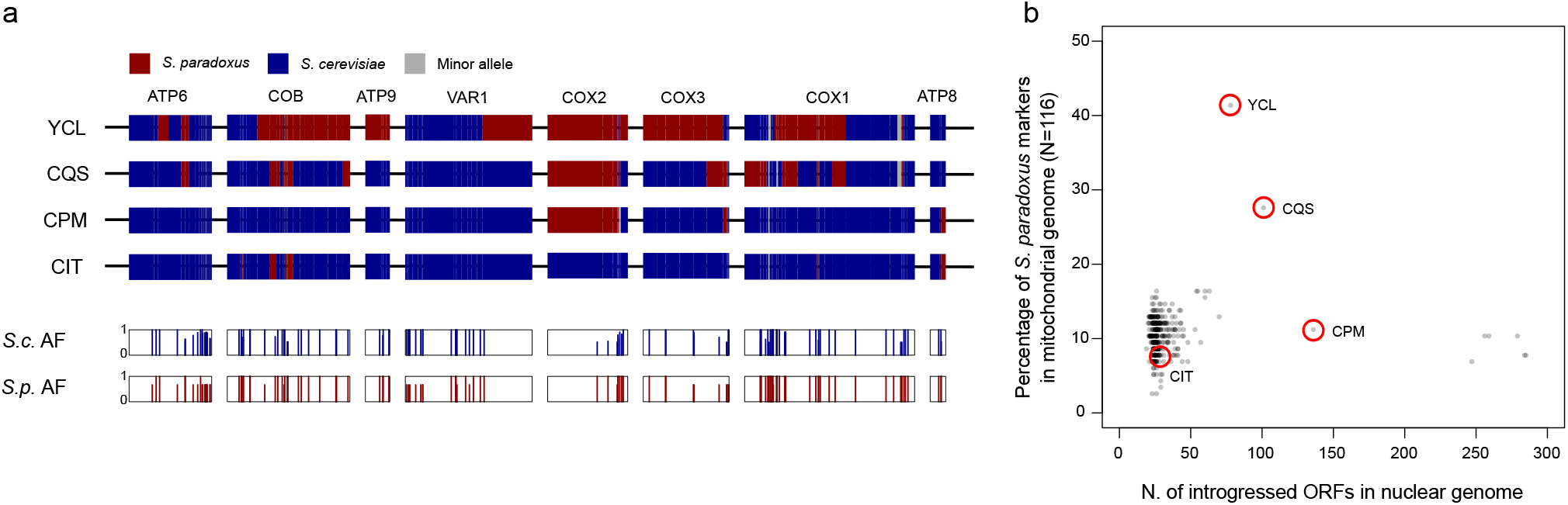
Rare *S. paradoxus* introgressions. (a) Polymorphic markers between *S. cerevisiae* and *S. paradoxus* across the mitochondrial CDSs were used to identify introgression events. Introgression boundaries are set as midpoint between markers. The two bottom rows indicate the frequency in the population of the major or consensus allele (AF), in the specific position and species. (b) Number of introgressed ORFs in the nuclear genome does not correlate with percentage of genetic markers of *S. paradoxus* in mitochondrial CDS. Only isolates with complete unambiguous CDS data were included (N=353). Position of isolate reported in panel (a) are circled in red.

We further extend the analysis of the 116 polymorphic sites to 353 isolates with fully assembled CDS. We observed additional potential cases of mitochondrial introgressions. The isolate YCL mitochondrial sequence harbours over 40%of *S. paradoxus* markers, possibly indicating a recombinant genome derived from a recent transfer event. In addition, a small number of *S. paradoxus* markers is found in each *S. cerevisiae* isolate, perhaps due to incomplete lineage sorting. Overall, the number of *S. paradoxus* markers in the mitochondrial genomes does not correlate with the number of introgressed ORFs in the nuclear genomes (fig. 3b), suggesting that the interspecies gene flows were independent due to distinct origin and/or fate.

### Introns gain and loss during evolution and dispersal

Two mitochondrial protein coding genes, *COB* and *COX1*, harbour introns at multiple sites and we explored their presence-absence patterns in the whole 1011 isolates collection. *COX1* introns are found at varying frequencies (median 0.48) with highly variable presence-absence profiles (fig. 4a). Introns patters further support low variability within North American, Malaysian and Mixed Origin lineages (supplementary fig. S5). In contrast, the groups of loosely related mosaics (M1, M2 and M3 clusters) show the lowest level of intron conservation, consistent with their admixed genetic backgrounds.

**FIG. 4.**
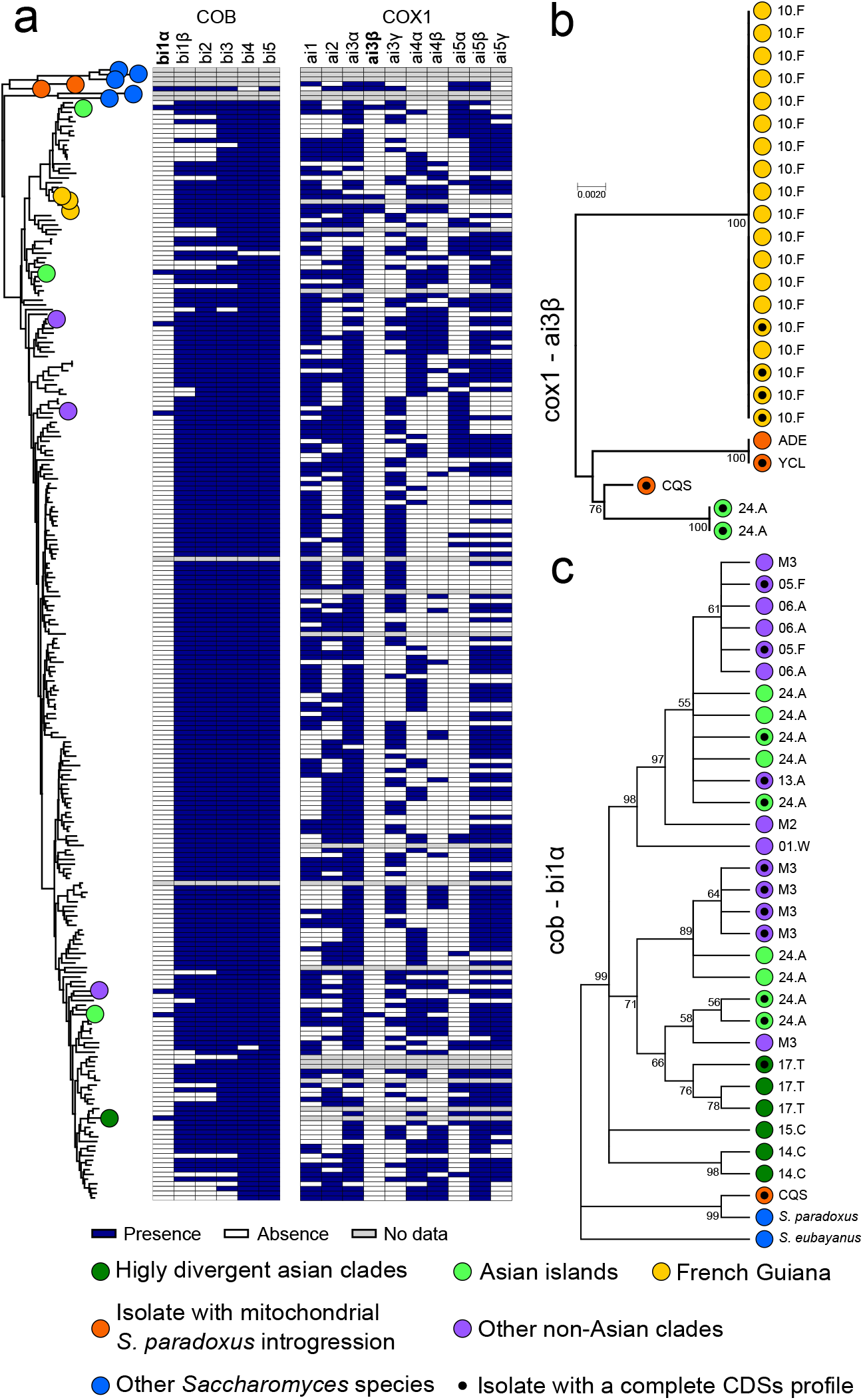
Introns phylogeny underlie both loss and gain events. (a) The distribution of introns presence and absence is not consistent with the mitochondrial tree phylogeny. The rare introns bi1α and ai3β are highlighted (bold), intron ai4γ was not found in the sequenced collection and not shown. (b) Only 4 non redundant sequences have been found for cox1 ai3β intron. Their sequences are unrelated to other *Saccharomyces* species, which could not be used for rooting. The peculiarity of the distribution of this intron could suggest a lineage-specific gain event. (c) Rooted tree of the *COB* intron bi1α using *S. paradoxus* and *S. eubayanus* sequences as out-group. Nodes with bootstrap values below 0.5 have been collapsed. Its presence in multiple highly divergent Asian lineages and in other *Saccharomyces* species is consistent with intron loss following the out-of-China dispersal. The isolate CQS, which harbours introgression in both nuclear and mitochondrial genome, also derive from S. paradoxus origin. This is compatible with the downstream exonic sequence, which also is introgressed.

The *COX1* introns frequencies in the population is consistent with previous report (Wolters et al. 2015), ranging from 26 to 86%. We identified a total of 103 different *COX1* intron combinations with two introns, ai4β and ai5α, that are never found together (ai4β is in 89 while ai5α in 85, out of of 408 alleles). Given the linkage between these closely spaced intronic positions, they either raised in two ancestral populations and are unlikely to be brought together by recombination or the double presence is functionally incompatible. Two additional *COX1* introns, ai3β and ai4γ, are very rare in *S. cerevisiae* population. While aI4γ is also absent in most related *Saccharomyces* species, ai3β intron is present in all of them. The only occurrence of ai3β previously reported in *S. cerevisiae* was in the YCL isolate, which also contains *S. paradoxus* introgression around the intron position in *COX1*. However, although the ai3β intron is present in *S. paradoxus*, the ai3β intron sequence of YCL is closer to the one found in *Lachancea meyersii* (Wolters et al. 2015). In addition to the YCL allele, we found other three variants of ai3β, all related to the *Lachancea* sequence. Two variants are present in YCL and ADE isolates with abundant *S. paradoxus* introgressions, while the CQS strain has related version. Additional ai3β intron are present in two Asian isolates and in 19 French Guiana isolates, whose clade is highly introgressed from *S. paradoxus* (fig. 4b). The presence of the ai3β intron among these highly introgressed lineages suggests separate lateral transfer events from *Lachancea*, although it cannot be ruled out that these introns where initially transferred from *Lachancea*, or a related genus, to *S. paradoxus* before the introgression occurred.

In contrast, the six *COB* introns a more uniformly present (frequencies ranging from 88 to 99%, fig. 4a) with the only exception of the recently described bi1α (Wolters et al. 2015) occurring at low frequency (~5%). Surprisingly, bi1α is common among the early-branching Asian clades (Peter et al. 2018). Other isolates harbour it, mainly mosaic isolates, but segregate at low frequency in non-Asian clades. Its presence in several *Saccharomyces* outgroup species and in the *S. cerevisiae* basal lineages suggests a loss preceding or during the out-of-Asia dispersal. The intron could have been introduced again, from secondary contacts with bi1α-positive Asian lines. To test these hypotheses, we constructed a phylogenetic tree using all the bI1α intron sequences and outgroups (fig. 4c). The bi1α phylogenetic tree shows more variants of Asian sequences compared to non-Asian ones, which mainly cluster in two groups stemming from separated branches of Asian introns, consistent with multiple separate regain events in the worldwide population.

### Structural rearrangements are rare in mitochondrial genomes

Next, we investigate the size and the presence of structural variation across the mitochondrial genomes. Considering the 250 circularized assemblies, the mitochondrial genome sizes range from 73,450 to 95,658 bp (supplementary table S3). As the gene content is entirely conserved between these isolates, this high size plasticity is driven by variability of the intergenic region (ranging from 45,254 bp to 69,807 bp), and the intron content (ranging from 7,748 bp to 20,024 bp in size) (supplementary fig. S6). Both factors are highly correlated to the total mitochondrial genome length (r^2^ 0.769 and 0.756, respectively) (supplementary fig. S7). Mitochondrial genome size is variable among isolates of the same lineage.

Synteny analysis across the 553 isolates with genome on single scaffold highlights four distinct genomic inversions (fig. 5, supplementary fig. S8). Two strains from the Wine/European and Ale beer lineages, BKI and AQT, share an inversion of the region that ranges from trnW to the *COX2* gene, while three closely related Wine/European strains (AIM, BNG and CFB) share a larger inversion that also encompasses the 15S rRNA gene (fig. 5b and 5c). Inversions were also found in BDN (African beer) and CDN (Ecuadorean) and are related to regions ranging from the 15S rRNA or *COX1* genes, respectively, to the *ATP6* gene (fig. 5d and 5e). All inversions boundaries map to the highly repetitive AT-rich intergenic regions, which prevents their precise delimitation. Interestingly, all these inversions lead to the loss of a feature shared by most ascomycetous yeast, namely that all mitochondrial protein-coding genes are transcribed from the same DNA strand (Talla et al. 2005). However, mitochondrial functions seem not to be impaired, as these isolates maintain their respiration capabilities.

**FIG. 5.**
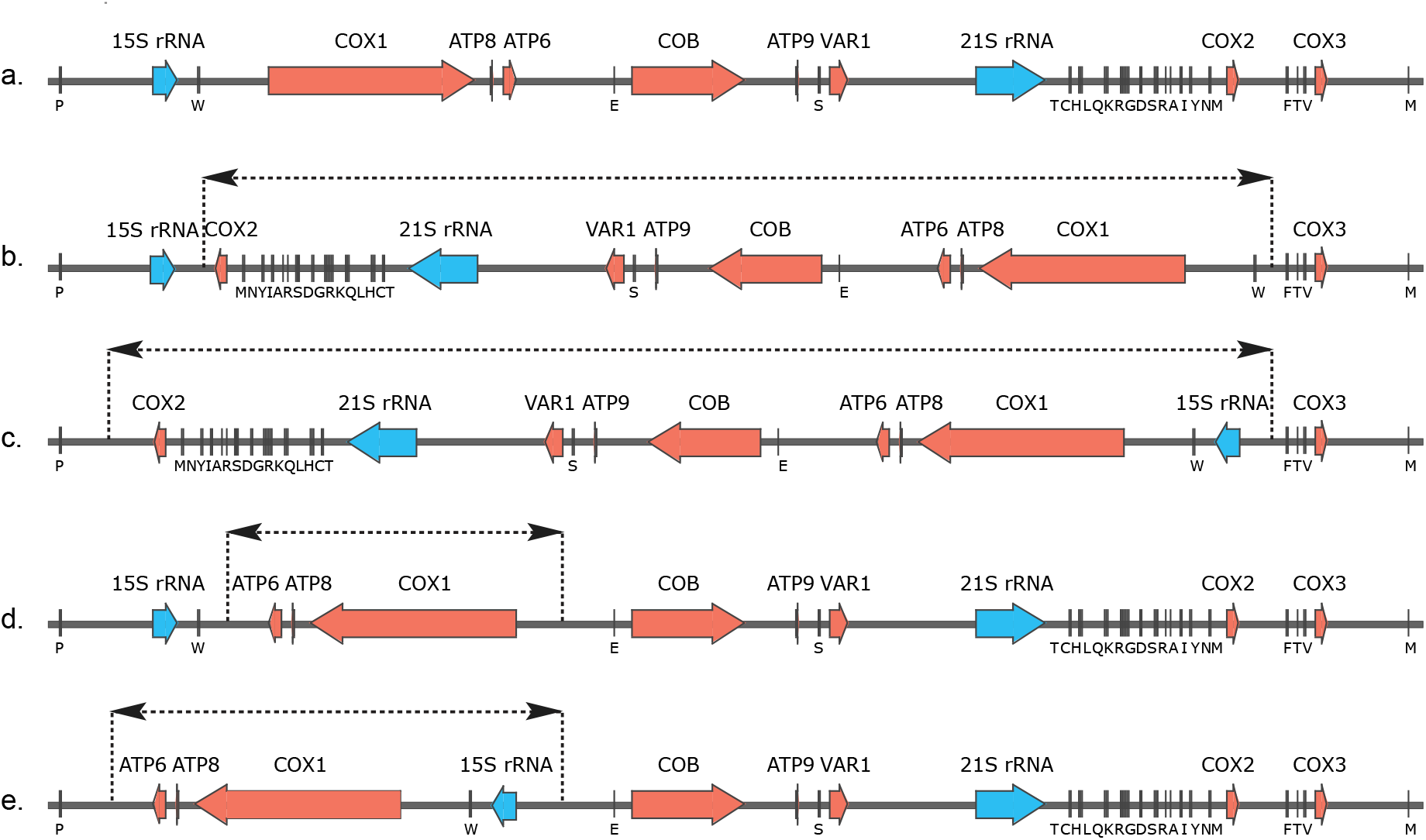
Structural variants in the mitochondrial genomes. Schematic of the mitochondrial genome organization annotated for protein-coding genes, rRNA and tRNA genes. The approximate breakpoint locations of the inversions are indicated by dotted lines. These mitochondrial genome organizations are related to different isolates: (a) S288C (shared by the vast majority of isolates), (b) AQI and BKI, (c) AIM, BNG and CFB, (d) CDN and (e) BDN.

Recent report suggested that the alteration of the gene order within yeast genera could be related to the mitochondrial genome size (Sulo et al. 2017). While the *Lachancea* and *Yarrowia* clades, with mitochondrial genome less than 50 kb, show high synteny across species (Friedrich et al. 2012; Gaillardin et al. 2012), the *Saccharomyces* clade (mitochondrial genome size > 65 kb) is more prone to rearrangements (Sulo et al. 2017). Indeed structural rearrangements were also detected in the mitochondrial genome of *S. paradoxus* (Yue et al. 2017). Our results suggest that mtDNA structural variation can be tolerated, perhaps restricted to balanced events that do not alter the CDS copy number.

### Variation in mtDNA copy number reveal natural petite isolates

Mitochondrial copy number can dramatically affect phenotypes but is hard to measure with high-throughput methods. We estimated mtDNA copy number using the relative coverage of *ATP6*, *COX2* and *COX3*, whose greater length and lack of introns give reliable mapping. The number of mitochondria is generally constant across clades (supplementary fig. S9), with no significant differences between domesticated and wild lineages, with a median of 18 mitochondrial genomes for each haploid nuclear genome. The variation is however particularly high across the population, reaching over 80 copies. The mitochondrial copy number scales up with ploidy in a linear way, with diploid strains having around double number of mitochondria and triploid having three times the number of mitochondria compared to haploid cells (fig. 6a). We run genome-wide associations to identify possible copy number genetic determinants, but no significant hits were detected in neither the nuclear and mitochondrial genomes.

**FIG. 6.**
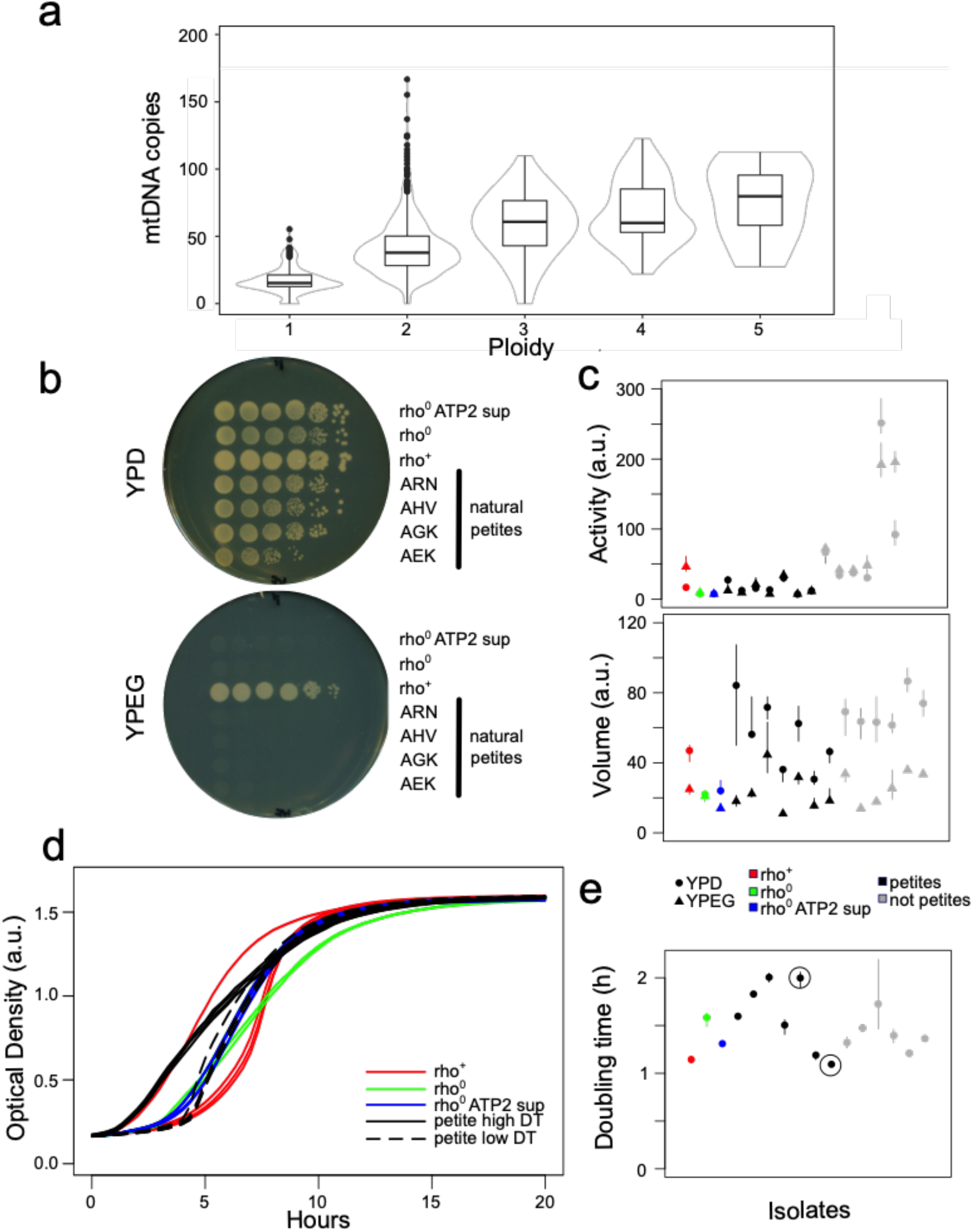
Natural variation in mitochondrial genome copy number. (a) Mitochondrial genome copy number linearly increase with the nuclear genome content. Fifteen natural petite isolates were detected. (b) Spotting assay on non-fermentable carbon source (YPEG) confirms natural petite isolates (a subset of tested isolates is shown). (c) Mitochondrial activity (as membrane potential) is strongly altered by lack of mitochondrial genome (black versus grey symbols), while the volume remains unaltered. (d) Growth curves variation of isogenic strains with normal mitochondria (rho^+^, red), petite (rho^0^, green) and petite harbouring the ATP2G1099T suppressing mutation (rho^0^ ATP2 sup, blue). Among natural petites, we can identify both isolates with high doubling time (DT, black solid line) and isolates with recovered growth rate, comparable to petite with suppressor mutations (black dashed line). (e) Generation times for isolates with different mitochondrial CN show at least two natural petite isolates that seem to have recovered normal growth rate on rich media. Growth curves for the circled isolates are shown in panel d.

Presence of mitochondrial genomes is assumed to be the natural state of *S. cerevisiae* cells, which is defined as rho^+^. However, strains can lose mitochondrial functionality under different conditions either by accumulating mutations (rho^-^) or by complete loss (rho^0^) of the mitochondrial genome. These mutants are defined as “cytoplasmic petite” (*i.e*. “small”) because they form small colonies in fermentation media due to their slow growth. Since the respiration-mediated ATP production is impaired in petite strains, they cannot grow in non-fermentable carbon sources. We identified 15 potential natural petites isolates (supplementary table S1) from coverage analysis and confirmed that they were unable to grow in media based on non-fermentable carbon sources (fig. 6b). Seven of them were laboratory-derived haploid (*HO* deleted) and since the manipulation could have caused their mitochondrial condition, they were excluded from further analyses. We examined a selection of four strains for mitochondrial activity (measured as membrane potential) and volume. Three strains were included as control, a wildtype rho^+^ strain and two derived rho^0^ variants with one wearing an additional mutation (*ATP2* G1099T), which partially restore growth in rich media (Michael Breitenbach, unpublished data). As expected, activity data show inability to grow on non-fermentable carbon sources (YPEG). There was, however, no significant variation between mitochondrial volume of wild type and petites isolates, consistent with the essentiality of maintaining mitochondria also in petite strains (fig. 6c, supplementary table S4). We investigated if these natural petites have fitness defect by running growth curves in rich media (YPD). The petite strains showed different growth rates, with two of them having normal growth fitness (fig. 6d and 6e, supplementary table S4). These strains do not have neither *ATP2* G1099T nor *ATP3* G348T polymorphism (also able to partially restore growth in rich media, Michael Breitenbach, unpublished data), hence other compensatory mutations might have restored fitness in these strains. Sporulation is known to be impaired in petite isolates (Briza et al. 2002) and consistently all natural petite isolates do not sporulate.

## Conclusions

The *S. cerevisiae* yeast has long been considered as an organism with an almost clonal reproduction and strictly uniparental mitochondrial inheritance (Dujon 2019). Nevertheless, the scenario emerging from our phylogenetic analyses revealed that outbreeding and recombination drive mitochondrial genome evolution. Quantitative variation between mitochondrial and nuclear genomes in admixture, population structures and sequence diversity underlie how differences in biology and selection of two genomes coexisting in the same cells can lead to highly discordant evolutionary trajectories. Furthermore, mitochondrial genomes are more refractory to interspecies introgressions, with only rare examples in the population. However, these rare events further support contribution from both parental strains to the mitochondrial sequence, implying transient coexistence of unknown timespan. This situation can lead to incomplete lineage sorting, as suggested by the allele frequencies distribution in *S. cerevisiae* and *S. paradoxus* mitochondrial genomes.

We observed a broad distribution of mitochondrial genome copy number across the population, of similar magnitude of variation generated in the knockout collection (Puddu et al. 2019), which broadly scales up linearly with nuclear genome ploidy. The occurrence of natural petite strains, remarkably able to growth in fermentation media without detectable defects, demonstrates how variation in the nuclear genome can compensate the complete loss of mitochondrial genomes. Overall, these observations further reinforce the strong functional interplay between the two coexisting genomes despite their discordant evolutive trajectories. Future efforts will reveal the molecular details of such two-genome interactions and quantify their contribution toward the species phenotypic variation.

## Acknowledgements

We thank Jozef Nosek for helpful discussions. This work was supported by France Génomique (ANR-10-INBS-09-08). GL is supported by Agence Nationale de la Recherche (ANR-11-LABX-0028-01, ANR-13-BSV6-0006-01, ANR-15-IDEX-01, ANR-16-CE12-0019 and ANR-18-CE12-0004). J.S. is supported by a European Research Council (ERC) Consolidator grant (772505). J.S. is a Fellow of the University of Strasbourg Institute for Advanced Study (USIAS) and a member of the Institut Universitaire de France.

## Material and Methods

### Mitochondrial genome assembly

We investigated a total of 1,011 *S. cerevisiae* isolates from the 1002 Yeast Genome Project (Peter et al. 2018). Among these, 84 isolates already had their complete mitochondrial sequence available (Strope et al. 2015). For all other isolates, we identified mitochondrial scaffolds from whole genome assemblies through similarity searches with the BLAST suite of programs (Altschul et al. 1997), using the S288C mitochondrial sequence (EMBL: KP263414) as a query. In total, 905 mitochondrial assemblies could be retrieved, consisting of 1 to 27 scaffolds (supplementary table S5). Among them, 468 mitochondrial assemblies consist of a single scaffold longer than 70 kb and 166 could be circularized by circulator (Hunt et al. 2015) bringing the total number of circularized assemblies to 250. The final assemblies were linearized so that their start corresponds to the starting position of the S288C reference genome (supplementary table S3). As results, 698 assemblies had the full set of CDSs available, 353 of which without any ambiguous position and were hence used for the phylogenetic inferences.

### Genes and coding sequences

The sequences of the eight protein-coding genes (*ATP6*, *ATP8*, *ATP9*, *COB*, *COX1*, *COX2*, *COX3*, and *VAR1*) were retrieved from the 1,011 assemblies by performing BLAST similarity searches. The boundaries of the coding regions of *COX1* and *COB* genes were identified through pairwise global alignments between the gene sequences and the reference CDS, constructed with stretcher (Myers and Miller 1988), and manually refined. The average pairwise diversity π was calculated with Variscan (Hutter et al. 2006) while sequence divergence was calculated as percentage of SNPs.

### *S. paradoxus* marker identification

Polymorphic positions were extracted by aligning the isolates CDSs to the S288C ones running MUMmer v3.0 (Kurtz et al. 2004) both for the *S. cerevisiae* collection and a set of PacBio sequenced *S. paradoxus* strains (Yue et al. 2017). The three American and Hawaiian *S. paradoxus* isolates were used while the European and Far East Asian isolates were removed due to their similarity to the *S. cerevisiae* mitochondrial sequence. SNPs were extracted with the command show-snps and the options –CrIT. For both species, each polymorphic position was evaluated to obtain the most common allele in the population, i.e. the major allele for *S. cerevisiae* and a consensus sequence for *S. paradoxus*. All positions where the major allele of *S. cerevisiae* and the consensus of *S. paradoxus* differ were retained to generate a marker list of 116 positions scattered across the genome. It is important to notice that, probably due to incomplete lineage sorting, in most of the cases the less common allele of one of the two species corresponds to the more common allele of the other one, which introduces a certain amount of noise among the selected markers.

### Population structure

A non-redundant database of profiles of mitochondrial CDSs was built for the isolates with complete CDS sequences. Variant Call Format (VCF) file was created running snp-sites v2.3.3 (Page et al. 2016) on the concatenation of the CDSs genes multiple alignment. Plink v1.90 (Purcell et al. 2007) was used to prepare the data to run ADMIXTURE v1.3.0 (Alexander et al. 2009) software. The concatenations of CDS were also used to run SplitsTree v4.16.6 (Huson and Bryant 2006) and produce the phylogenetic network while the bootstrap of the NJ phylogenetic tree was produced by MEGA v7.0.26 (Kumar et al. 2016).

### Identification of rearrangements

The gene coordinates were determined on the one-scaffold assemblies by running tRNAscan for tRNA and BlastN similarity searches for rRNA and protein coding genes. Structural rearrangements were than detected by loss of synteny. The rearranged assemblies were further investigated through alignment with the reference mitochondrial genome with MUMmer v3.0 (Kurtz et al. 2004), nucmer was used to align the sequences and plots were generated with mummerplot.

### Copy number estimation

The number of copies of the mitochondrial genome was assessed by mapping, using BWA v0.7.15 (Li and Durbin 2009) with the option –U 0. The reads of individual strains of three mitochondrial CDSs (*ATP6*, *COX2*, *COX3*) gave the most reliable copy number estimation. These CDS were chosen for lack of introns and their size. Samtools view was used to filter the results with the option –q 20. The copy number for haploid genome was estimated as the ratio between the average of the median coverage for the three mitochondrial CDS and the nuclear genome coverage (median of the median coverage for each chromosome). The estimated total copy number for each isolate was calculated as this copy number times the ploidy.

### Spotting assay

Cells were pre-grown overnight in liquid YPD before being diluted in water. For each 10-fold dilution, 5 μL were spotted either on YPD or YPEG (3% ethanol, 3% glycerol, 1% yeast extract, 2% peptones) agar plates. Plates were then incubated 48 hours at 30 °C and images were acquired with a Epson Perfection V330 scanner.

### Growth curves

A YPD overnight culture was 1:100 diluted into fresh YPD in a 96 well plate. Cells were incubated for 48 hours at 30 °C in a plate reader (Tecan, Infinite F200 Pro). Optical density (600nm) was monitored every 20 minutes. Four replicates were run for each strain. PRECOG software (Fernandez-Ricaud et al. 2016) was used to calculate the doubling times.

### Mitochondrial activity and volume

Cells grown overnight in liquid YPD were diluted 40X in 200 μL of fresh YPD or fresh YPEG (1% yeast extract, 2% peptones, 3% ethanol, 3% glycerol) in a 96 well plate and incubated 5 hours at 30°C. Respectively 8 μL/30 μL of cells from YPD/YPEG were then transferred into 192 μL/170 μL of a HEPES 10mM – glucose 2% solution. Samples were washed twice with HEPES-glucose and finally resuspended in 100 μL of mitochondrial staining solution (HEPES-Glucose + 100nM MitoTracker Green (Molecular Probes) + 20 nM MitoTracker Deep Red (Molecular Probes)) in a 96 well plate and incubated 30 min in the dark at 30 °C. MitoTracker Green is known to accumulate passively into mitochondria proportionally to their volume, whereas MitoTracker Deep Red is known to accumulate into mitochondria proportionally to their membrane potential. The samples were analysed on a FACS-Calibur using the HTS module. The relative mitochondrial volume was estimated by reading fluorescence in the FL-1 channel, whereas the relative mitochondrial activity was estimated by reading fluorescence in the FL-4 channel.

### Petite isolate sporulation activity

The 1,011 sequenced collection was phenotyped for sporulation efficiency (De Chiara *et al*, in preparation). Isolates were pre-cultivated in yeast peptone dextrose (YPD; 2% dextrose, 1% yeast extract, 2% peptone, 2% agar) before being diluted 1:50 into 10 mL of pre-sporulation media (YPA; 2% potassium acetate, 1% yeast extract, 2% peptone) and grown 48 hours at 30 °C (shaking = 250 rpm). Pre-sporulated cells were transferred to sporulation media (2% potassium acetate) into 250 mL flasks at a 5:1 volume/medium ratio and a final optical density (OD600) of 0.6. Flasks were kept at 23 °C and shaken at 250 rpm. To estimate sporulation efficiency, >100 cells/sample were counted at 24- and 72-hours post-transfer to sporulation medium using an optical microscope (Zeiss Axio Lab.A1). Sporulation efficiency was estimated as the number of asci divided by the total cell count.

## Supplementary tables

Table S1 Overview of the *S. cerevisiae* collection

Table S2 Genetic diversity metrics within the population and per subpopulation

Table S3 Overview of the complete mitochondrial genome assemblies

Table S4 Phenotypes of the petite isolates

Table S5 Topology of all mitochondrial genome assemblies

## Supplementary data

Within the 1002 Yeast Genome website, we provide a dedicated directory (http://1002genomes.ustrasbg.fr/files/MitochondrialGenomes) that provide access to:

- mitochondrialAssemblies.tar.gz: mitochondrial assembly for each isolate (.fasta format per isolate).
- allGeneSequences.tar.gz: sequences of all complete CDS and both CDS and gene for COX1 and COB (a.fasta format per gene/CDS).
- allNonRedundantAlleles.tar.gz: all the alleles and allele combinations considered within our population (.fasta format per gene + xlsx summary file)

