## Supplemental figures for "Discordant evolution of mitochondrial and nuclear yeast genomes at population level"

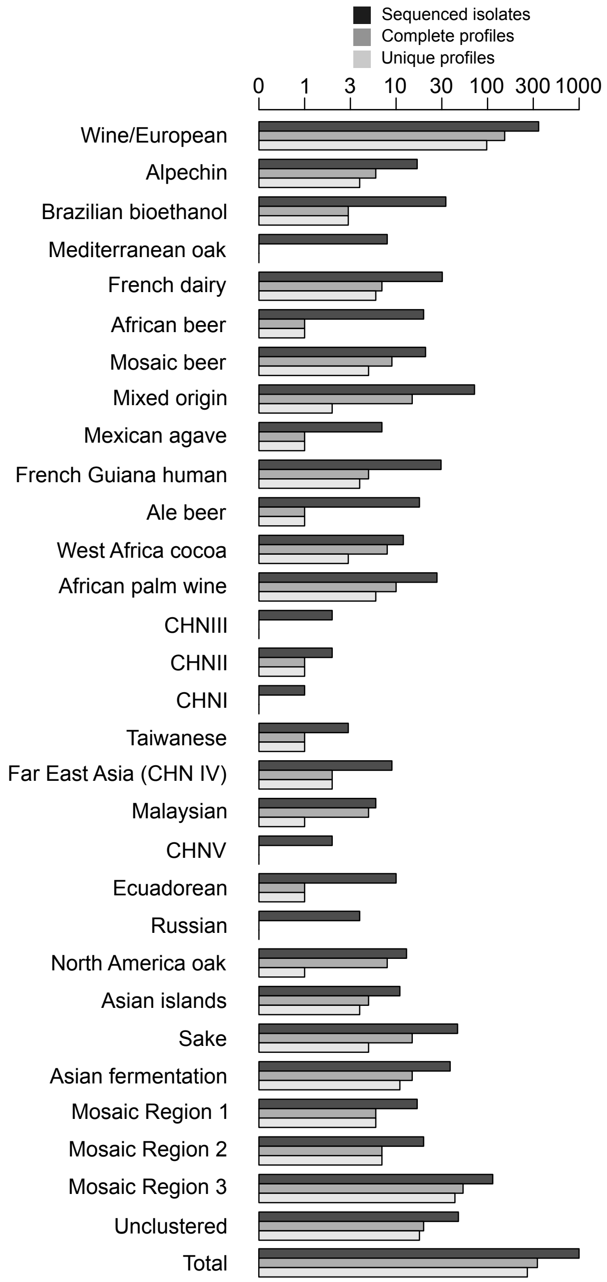


**Fig. S1 Dataset overview**

Dataset overview for each clade (named as in Peter et al. 2018) for number of isolates with genome sequenced, complete CDSs assembled and non-redundant profiles.

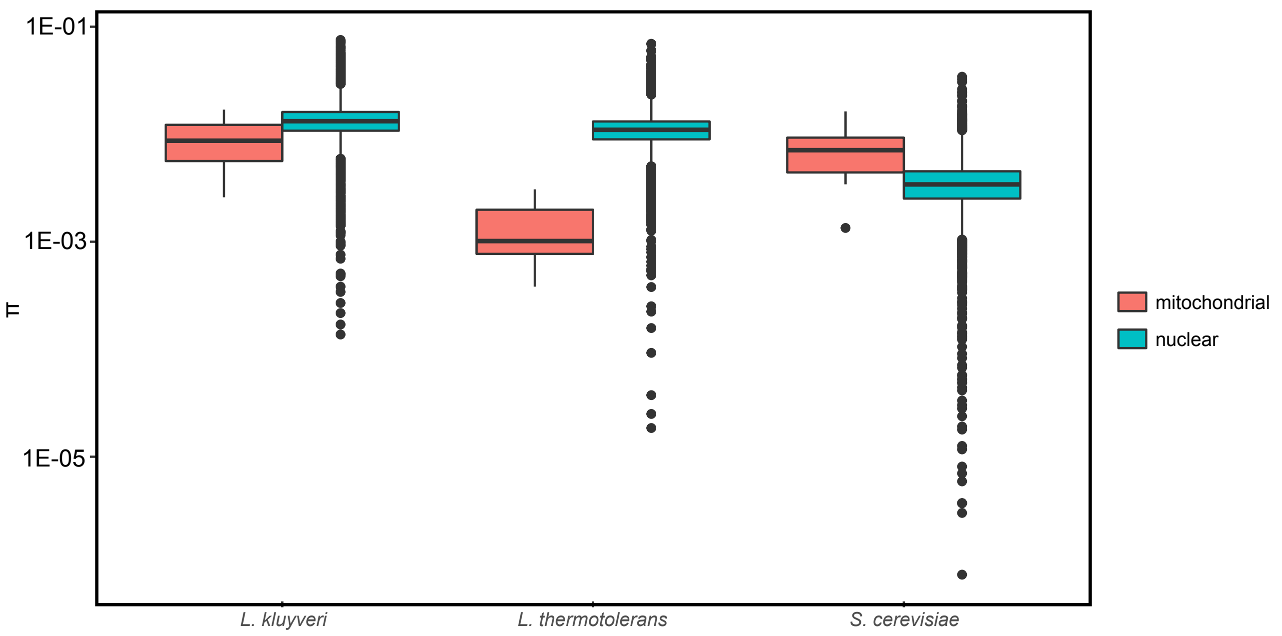


**Fig. S2 Genetic diversity of nuclear and mitochondrial genes for three yeast species**

Distribution of the π values of mitochondrial and nuclear protein coding genes for *S. cerevisiae* and two other yeast species, *Lachancea kluyveri* and *Lachancea thermotolerans*. *S. cerevisiae* is the only species for which genetic diversity is higher in mitochondrial genome compared to nuclear genome.

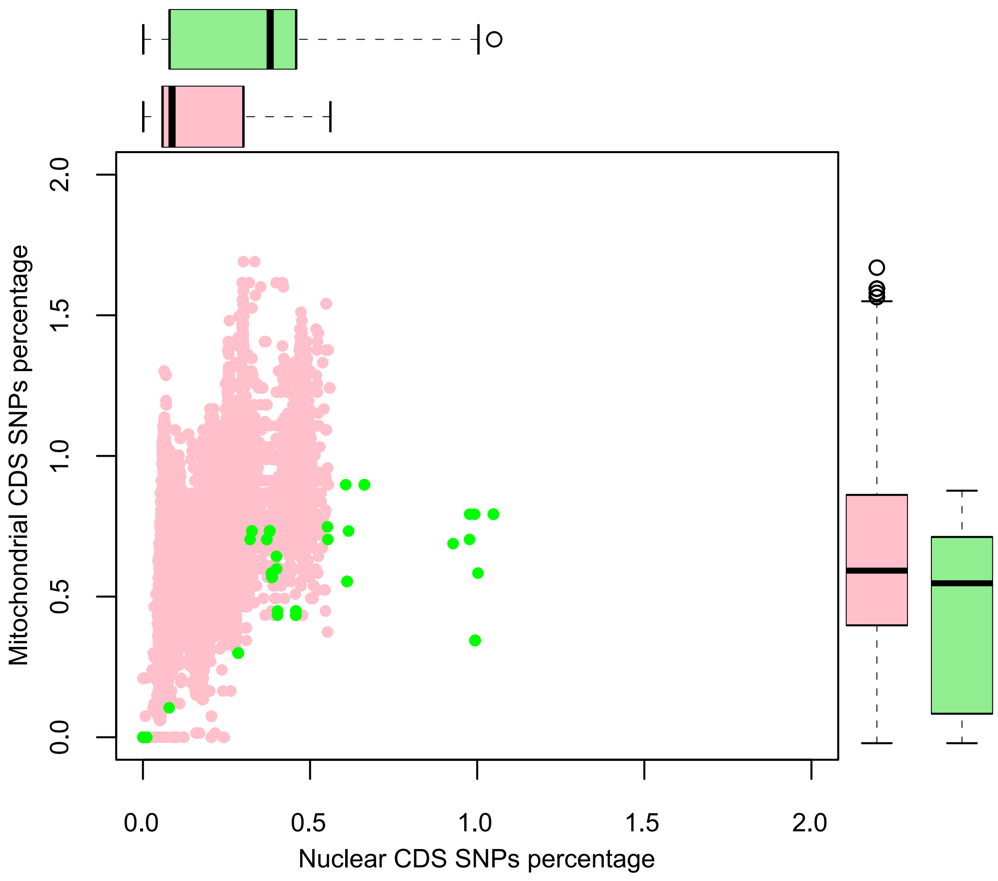


**Fig. S3 Inter-clade distances for domesticated and wild lineages**
Pink dots represent distances between isolates belonging to domesticated clades, green dots represent distances between isolates belonging to wild clades. In wild clades, the mitochondrial differences do not scale up with the nuclear distance. In domesticate clades higher diversity is found for lower nuclear distances compared to the wild clades.

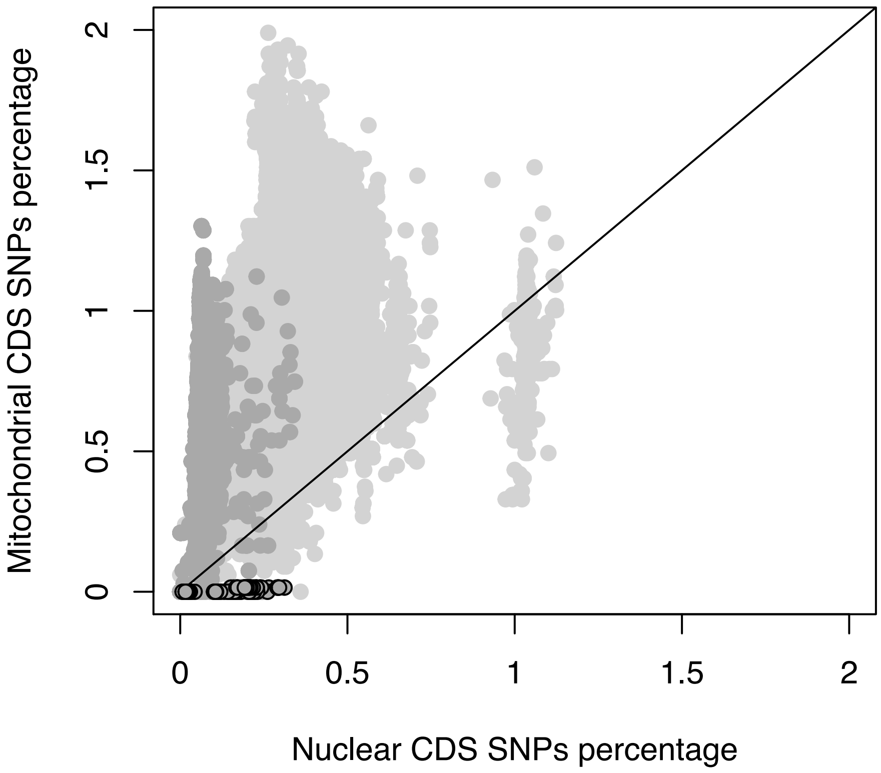


**Fig. S4 Scatterplot of intra-clades CDS SNPs percentages**

Light grey dots represent distances between isolates belonging to distinct clades while dark grey dots represent distances between isolates belonging to the same clade. Dark grey dots circled in black represent isolates belonging to the Mixed origin clade. The dashed line represents the equivalence between the two distances. Dots below the line represent isolate pairs whose mitochondrial distance is lower than the genomic distance. Mixed Origin clade have higher variation in genomic CDS than mitochondrial CDS. Only isolates with complete CDS data have been used (N=353).

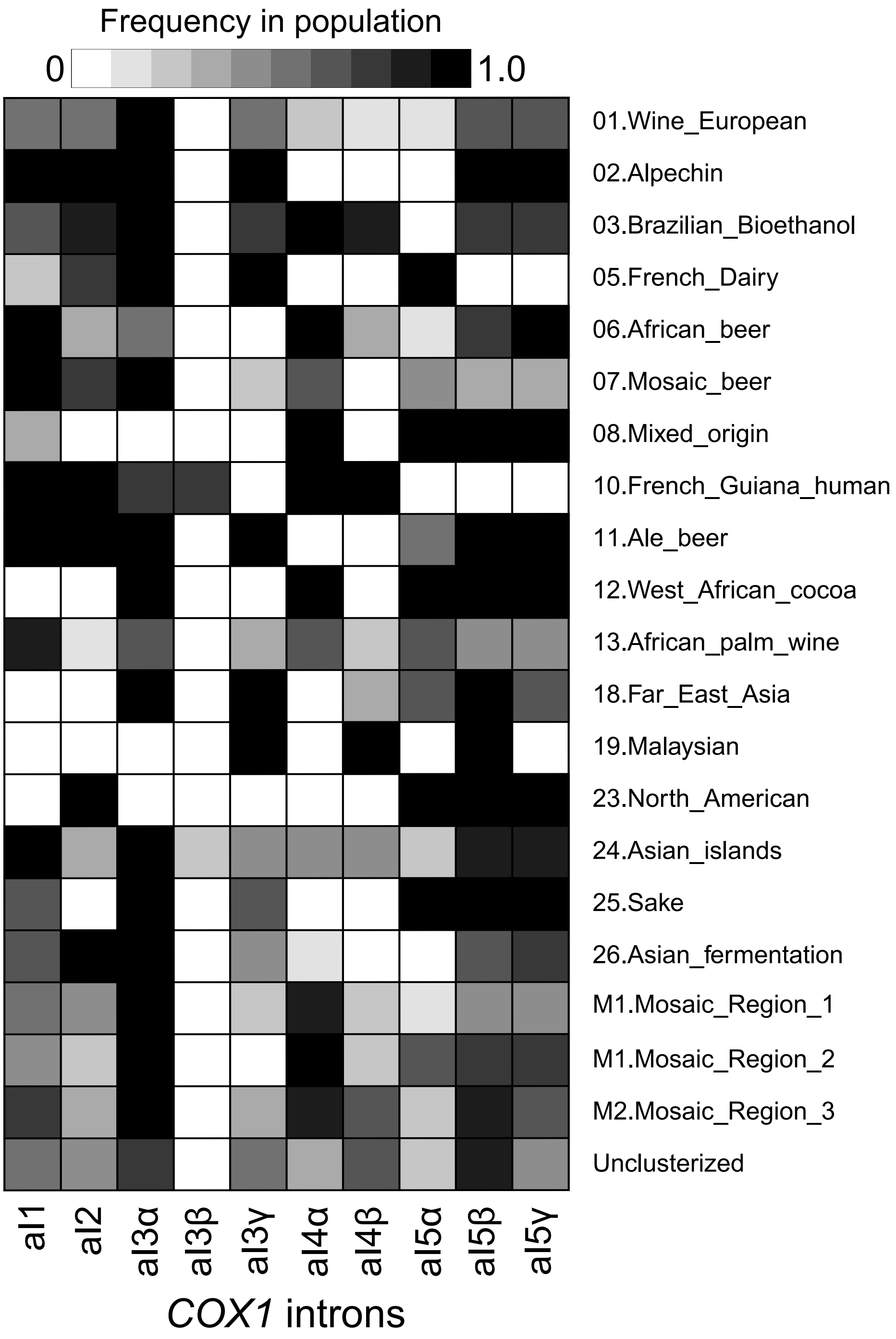


**Fig. S5 Frequency of different *COX1* introns**

The heatmap shows the frequency of the *COX1* introns across the *S. cerevisiae* nuclear clades. Black cells indicate presence in >90% the isolates, white cells indicate <10% in the clade.

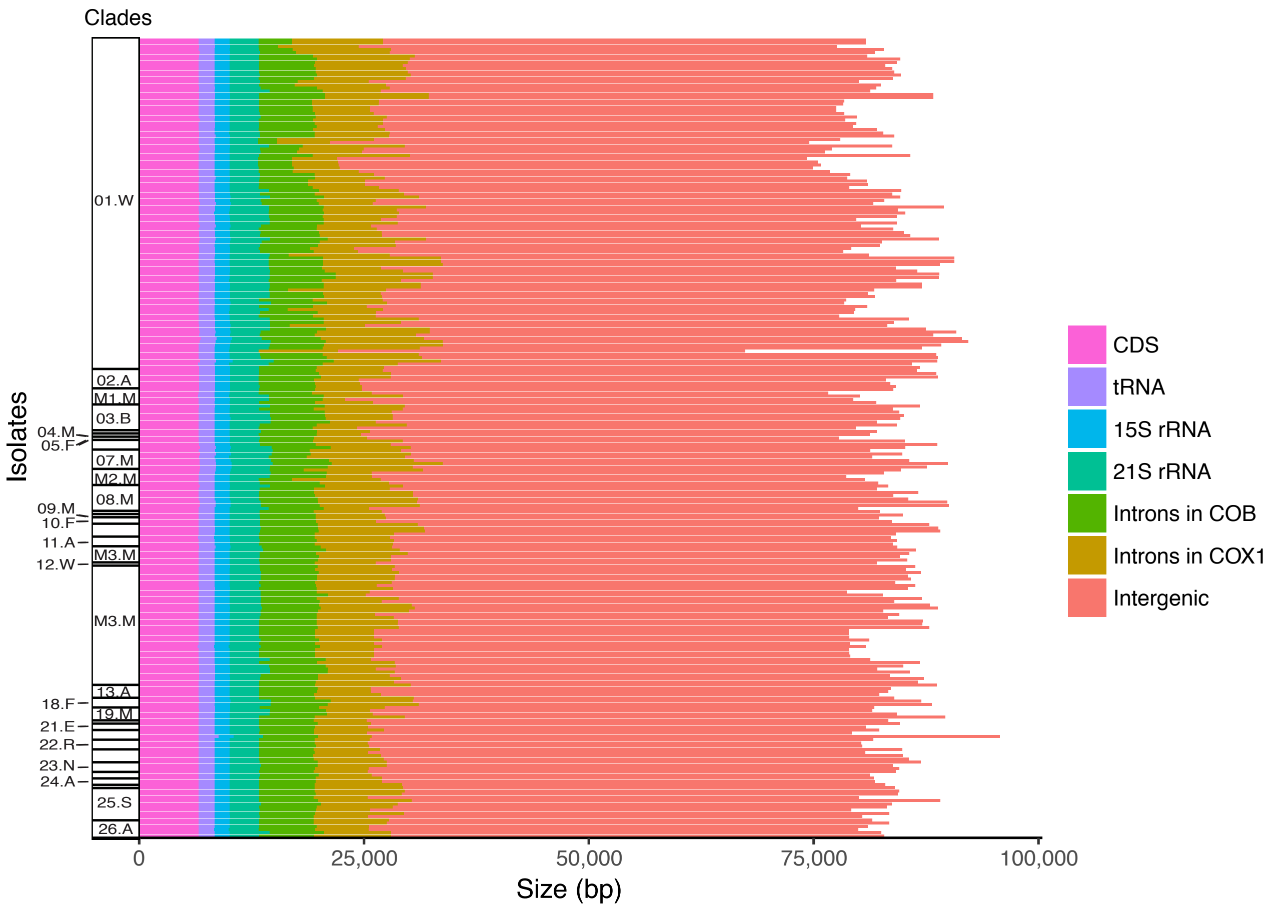


**Fig. S6 Mitochondrial genome size variation**

Size of all genetic elements located on the 250 circularized assemblies, grouped by clade (as described in Peter et al. 2018).

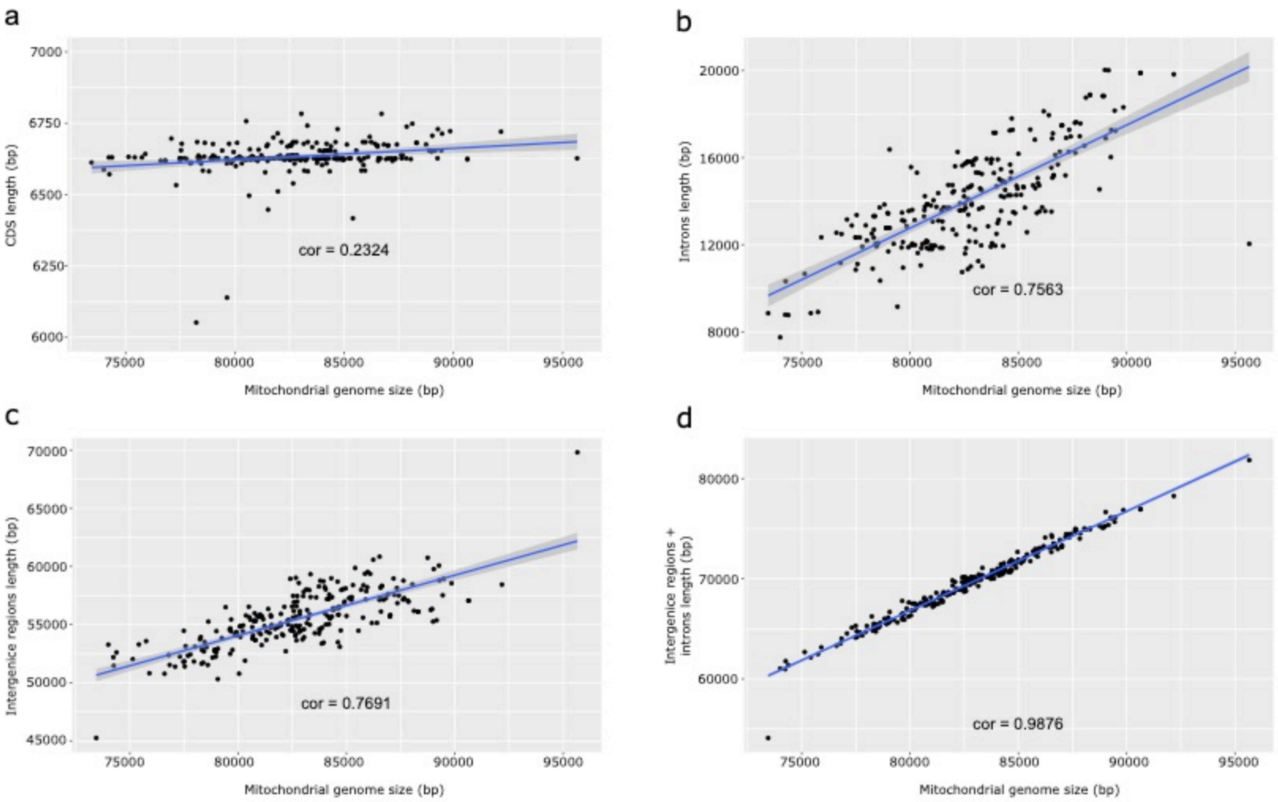


**Fig. S7 Mitochondrial genome size variation is driven by introns and intergenic regions**

Correlation between the length of the mt genome and the cumulative size of the (a) CDS, (b) introns, (c) intergenic regions, (d) intergenic regions and introns (p-values associated with correlation coefficients < 2.0E-04).

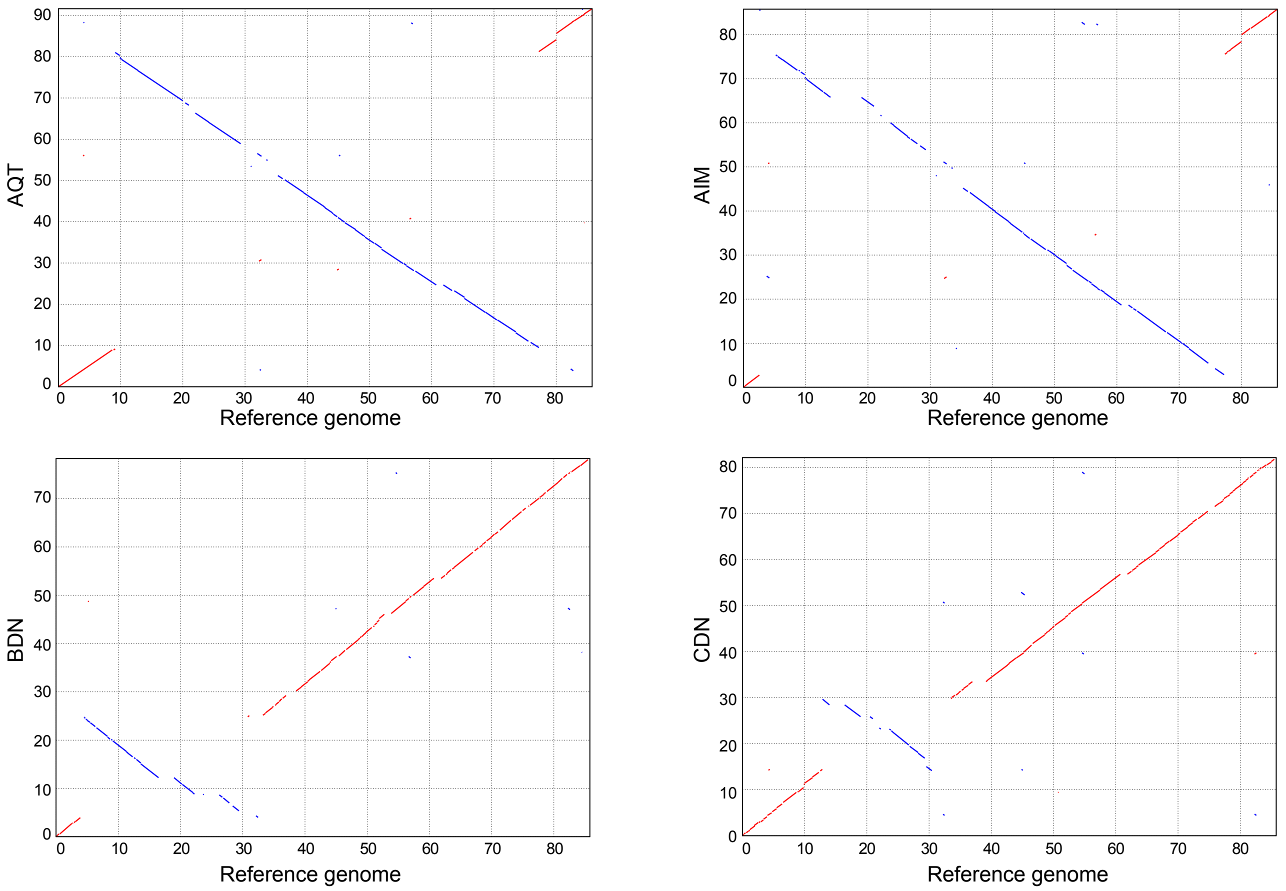


**Fig. S8 Structural variation in mitochondrial genomes**

Dotplots comparison between the reference genome and the mitochondrial assembly of 4 isolates showing different large inversions.

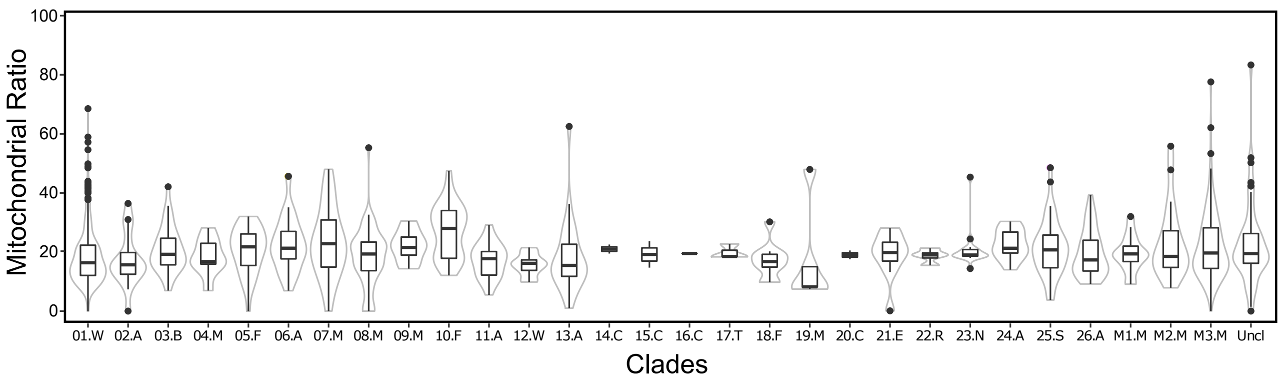


**Fig. S9 Copy number of mtDNA across clades**

The copy number is calculated as ratio with nuclear genome, to subtract variation derived by the ploidy. Mitochondrial genome copy number is relatively uniform (median ~ 18 copies) with variations in few clades *e.g.* increased copy number in French Guiana (10.F) isolates.
